# MicrobiomeDB: a systems biology platform for integrating, mining and analyzing microbiome experiments

**DOI:** 10.1101/176784

**Authors:** Francislon S. Oliveira, John Brestelli, Shon Cade, Jie Zheng, John Iodice, Steve Fischer, Cristina Aurrecoechea, Jessica C. Kissinger, Brian P. Brunk, Christian J. Stoeckert, Gabriel R. Fernandes, David S. Roos, Daniel P. Beiting

## Abstract

MicrobiomeDB (http://microbiomeDB.org) is a data discovery and analysis platform that empowers researchers to fully leverage experimental variables to interrogate microbiome datasets. MicrobiomeDB was developed in collaboration with the Eukaryotic Pathogens Bioinformatics Resource Center (http://EuPathDB.org) and leverages the infrastructure and user interface of EuPathDB, which allows users to construct *in silico* experiments using an intuitive graphical ‘strategy’ approach. The current release of the database integrates microbial census data with sample details for nearly 14,000 samples originating from human, animal and environmental sources, including over 9,000 samples from healthy human subjects in the Human Microbiome Project (http://portal.ihmpdcc.org/). Query results can be statistically analyzed and graphically visualized via interactive web applications launched directly in the browser, providing insight into microbial community diversity and allowing users to identify taxa associated with any experimental covariate.

## INTRODUCTION

Advances in high-throughput sequencing technology, together with the development of multiplex protocols for large-scale marker gene based studies (1, 2), have revolutionized microbiology, allowing scientists to complement culture-based approaches with culture-independent profiling of complex microbial communities (often referred to as a ‘microbiome’). As a result, there has been a tremendous increase in microbial census data generated from diverse habitats, including soil, ocean, the built environment, humans and animals. The growth of available data underscores a need for the development of web-based tools that allow users to rapidly explore public datasets, produce customizable visualizations, and generate hypotheses, without investing in compute resources or possessing extensive knowledge in bioinformatics or statistics.

Microbiome experiments are often accompanied by study designs that describe various attributes of the samples being studied. These ‘sample details’ can include information about the source from which the sample was derived, quantitative or qualitative biometrics from human or animal clinical studies, technical comments about how samples were processed and sequencing assays were carried out, respondent survey data, and much more. Despite the considerable effort made over the past decade to develop pipelines for analysis of raw sequence data generated from 16S rRNA marker gene studies (3, 4) and ‘shotgun’ metagenomic studies (5, 6), resources that allow scientists to interrogate the microbial community census data from the perspective of sample details are scarce. Web-based resources for microbiome researchers have focused largely on the storage and analysis of raw sequence data (7–9), or on visualization tools (10, 11), rather than on integrating data mining and analysis tools that include both sequence and sample-associated data. To address these unmet needs, we developed MicrobiomeDB (http://microbiomeDB.org) to empower users to identify experimental variables associated with changes in microbial community structure. We reasoned that the user interface and query tools developed by the Eukaryotic Pathogens Bioinformatics Resource Center, EuPathDB (http://EuPathDB.org) (12), for identifying genes of interest in eukaryotic pathogens based on gene attributes (*e.g.,* size in kilobases, expression level from RNA sequencing data, polymorphisms from DNA sequencing data) could be adapted to identify samples of interest from microbiome studies based on sample attributes (*e.g.,* age, sex, antibiotic exposure).

## DATA LOADING

A key feature of MicrobiomeDB is the development of an automated workflow for loading data from microbiome experiments (**Figure 1**). Microbial community census data from five publically available datasets **(Table S1)**, each as a Biological Observation Matrix (.biom) (13), was used as input for the workflow. Taxonomy assignments from the.biom file are mapped to the GreenGenes database (14, 15) to retrieve full 16S rRNA gene sequences, NCBI taxon identifiers, and taxonomy strings. Terms provided with the datasets to describe sample details included those compliant with the MixS (Minimal Information about any (x) Sequence) standard (16); however, most of the provided terms were not covered by the standard. Harmonization of sample details was performed to facilitate data integration and guide organization of sample descriptions for searches. Terms were mapped to the Open Biomedical Ontologies (OBO) Foundry ontologies (17) including the Environmental Ontology (18) and the Ontology for Biomedical Investigations (OBI) (19). The mapped ontology terms were added to the EuPath ontology, an application ontology developed for annotation of EuPathDB datasets (20). Unmapped terms were manually curated to generate new ontology terms that were then added. A web interface terminology was generated using a subset of the EuPath ontology. It contains only those terms needed for MicrobiomeDB and is used to guide searches based on sample details. Sample details were then formatted as an Investigation, Study, Assay file (ISA) (21) and, along with the census data, were structured using a Genomics Unified Schema, version 4 (GUS4; http://www.gusdb.org/SchemaBrowserBeta/categoryList.htm) (22) for loading into the database.

**Figure 1:**
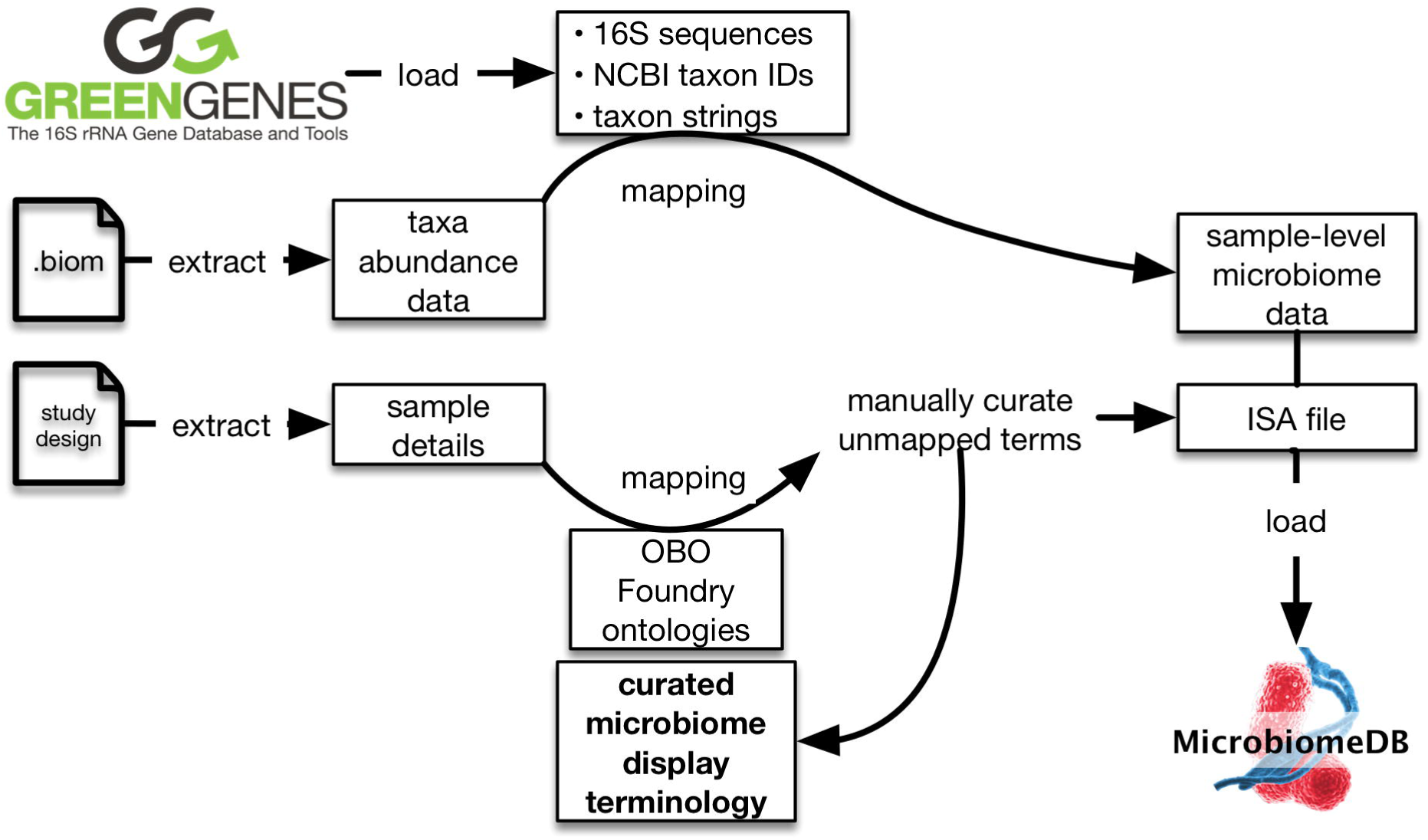
Schematic showing the automated data loading workflow for MicrobiomeDB. Greengenes identifiers are extracted from.biom files containing microbial community census data and used to retrieve NCBI taxon identifiers, full 16S rRNA gene sequences, and taxon strings. User-provided sample details are mapped to an OBO Foundry ontology and manual curation is used to produce a custom microbiome display terminology. Sample details are then formatted as an Investigation, Study, Assay (ISA) file and, along with microbiome census data, are structured in a GUS4 schema for loading into MicrobiomeDB.

## HOW TO USE MICROBIOMEDB

### The homepage

The homepage for MicrobiomeDB is divided into three main sections. The menu bar at the top of the page allows users to initiate a new search, access data sets, log in to view their saved searches, or contact our development team to report problems with the site or request new features or data to be added. The left-hand side bar displays social media content related to MicrobiomeDB, and provides users with access to the about page, as well as release notes and data sets. Finally, the main page summarizes the content of database, and provides access to tutorials and example searches.

### Performing a search in MicrobiomeDB

Searching the database is initiated by clicking on ‘New Search’ in the menu bar. Users have the option of searching MicrobiomeDB by either sample details or taxon abundance. Initiating a search by sample details takes users to a filter page (**Figure 2)** where all terms that describe all the samples in the database are available as a list (**Figure 2A**), searchable using a reactive text box (**Figure 2, arrow**). Selecting a term from this list displays its values and the number of samples that map to each value to the right (**Figure 2B**). When the user selects one or more values, the database is immediately filtered to return only samples annotated with the selected value(s). The user continues this process of filtering based on sample details, and can apply as many filters as they choose. Each filtering step produces a filter criteria that records and summarizes the user’s filter history, providing convenient, single-click access to return to and modify any prior filter step (**Figure 2C**).

**Figure 2:**
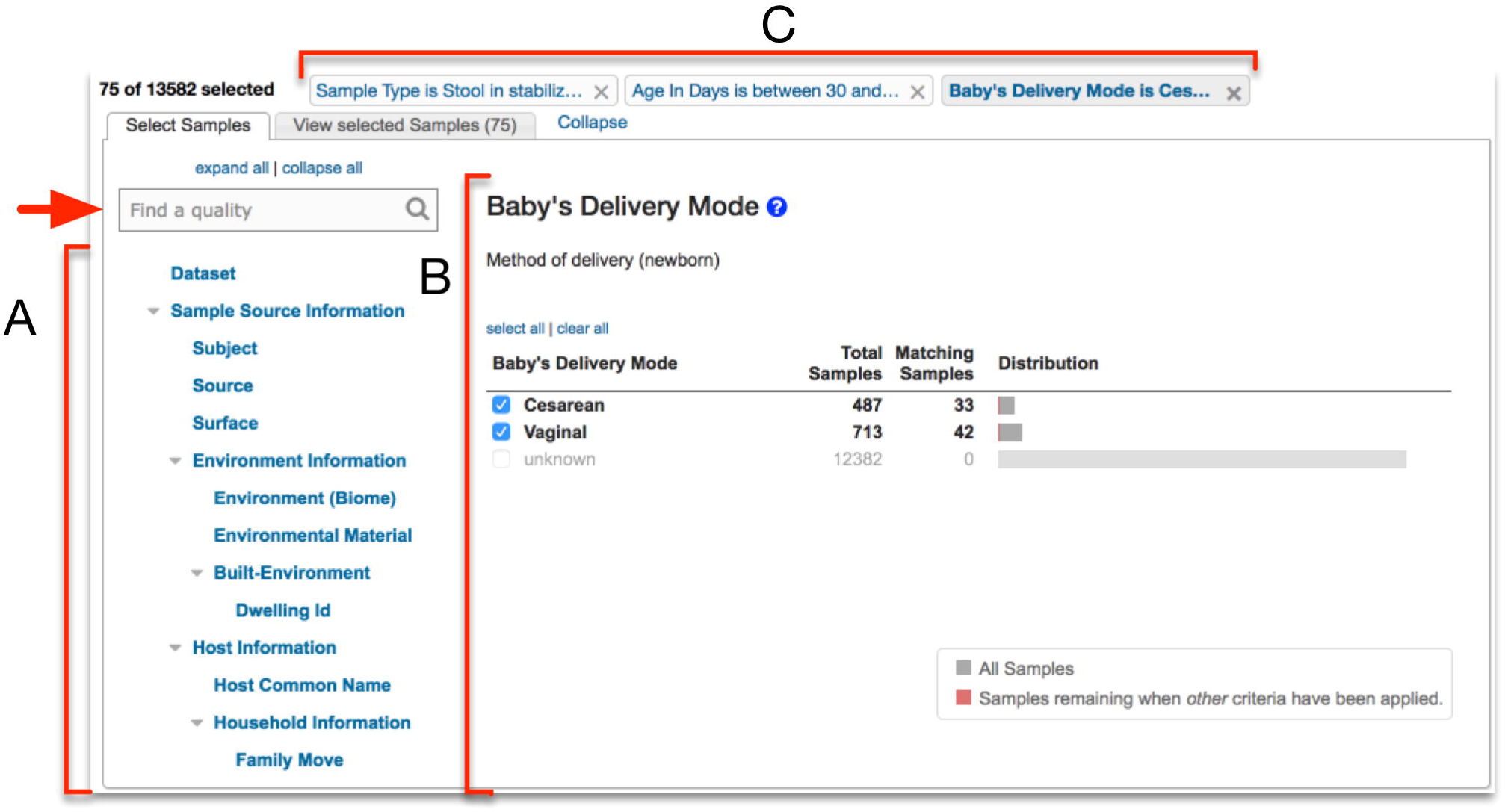
Screenshot of the filter page for searching by sample details. (A) The filter list shows all sample details describing all the samples in the database. This list is searchable via a reactive text box (red arrow) (B) Selecting any term from the filter list shows all the values associated with that term and the number of samples from the database that match each term. (B) Users filter the samples in the database by selecting values of interest. (C) Any filter applied by the user remains accessible through filter history at the top of the page.

A second way to search the database is by taxon abundance **(Figure 3)**, which also takes users to a filter page, but rather than using sample details to filter the database, they are presented with a list of the full taxonomy for all taxa represented in the database (**Figure 3A**). Searching for and selecting a single taxon from this list displays a distribution of the relative abundance for that specific taxon across all samples in the database **(Figure 3B)**. The user can select a single relative abundance value or a range of values by clicking and dragging any interval on the distribution **(Figure 3B, arrow)**, which then returns from the database only samples that contain the selected taxon at the specified relative abundance(s). Like the sample details filter page, the taxon abundance filter page records filter history (**Figure 3C**).

**Figure 3:**
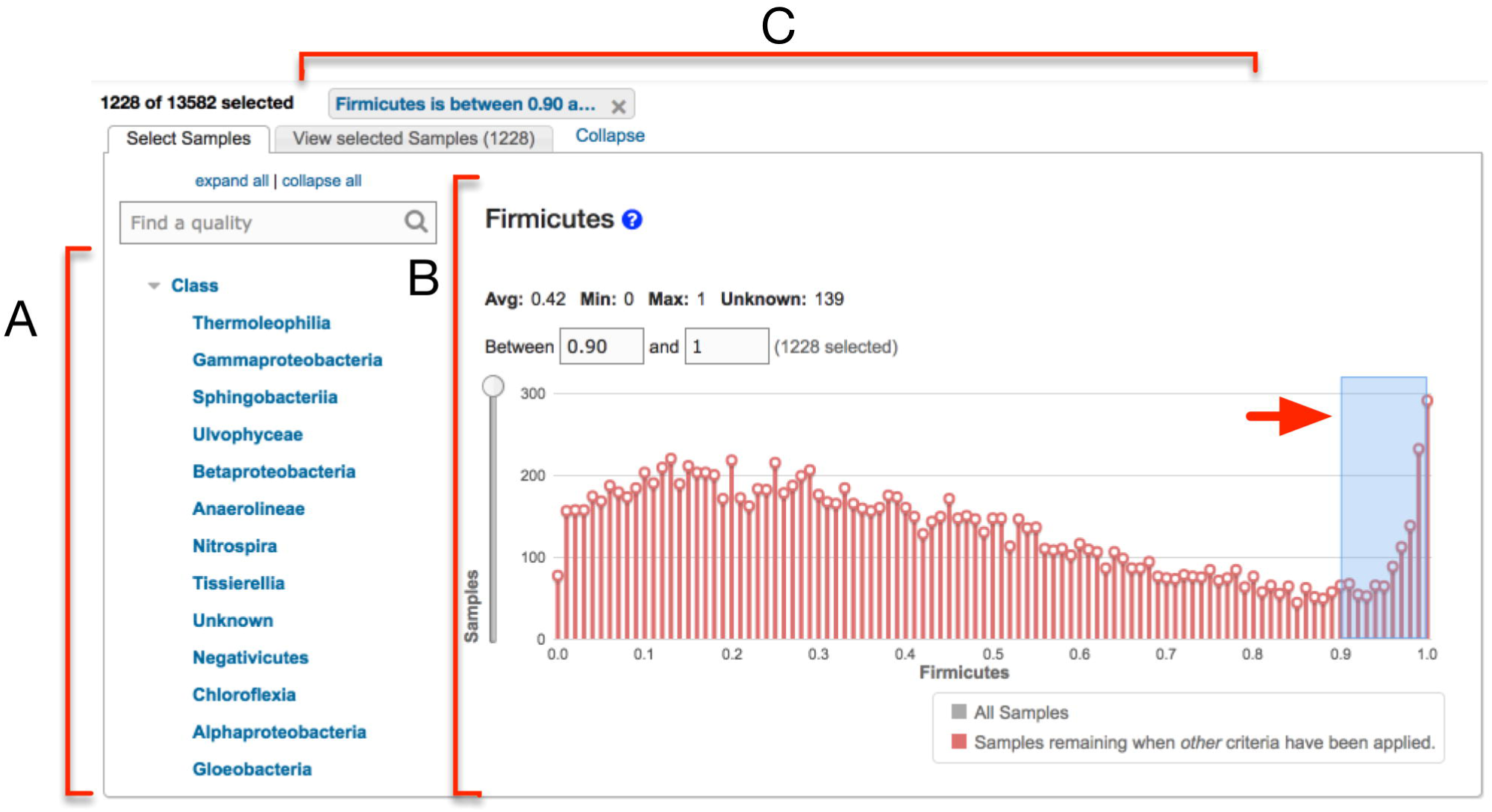
Screenshot of the filter page for searching by taxon abundance. (A) A searchable filter list shows all taxa at the phylum, class, family, order, genus and species level, across all samples in the database. (B) Selecting any taxon from the filter list shows the distribution of relative abundance for all samples in the database. Selecting any relative abundance value, or a range of values, applies a filter to the database that returns only samples that contain that taxon at the specified relative abundance value(s). (C) Any filter applied by the user remains accessible through filter history at the top of the page.

### Building a search strategy in MicrobiomeDB

Once filtering by sample details or taxon abundance is complete, selecting ‘Get Answer’ on the filter page takes the user to a results page **(Figure 4),** which is divided into two main panels. At the top of the page is the strategy panel (**Figure 4A**), where users can design *in silico* experiments by expanding on a single query to combine multiple queries of the database using Boolean operators. User-defined names and descriptions can be entered for each strategy, and strategies can be saved, kept private, made public, or shared with colleagues via a URL, making complex data mining strategies transparent and reproducible. Below the strategy panel is the sample table, which displays all the samples returned by the query (**Figure 4B**). These results can be downloaded (**Figure 4B, black arrow**), and sample-level details can be viewed by clicking on individual sample identifiers in this table, which takes the user to a sample record page where the dataset and any publications associated with the sample are listed, along with all sample details and taxon abundances recorded for the sample (**Figure 4C and 4D, respectively**). Taxon-specific details (*e.g.* available genome sequences) can also be accessed directly from the abundance table by clicking on the taxon identifier to navigate to either the National Center for Biotechnology Information (NCBI) Taxonomy Browser (23) or the Pathosystems Resource Integration Center (PATRIC) (24).

**Figure 4:**
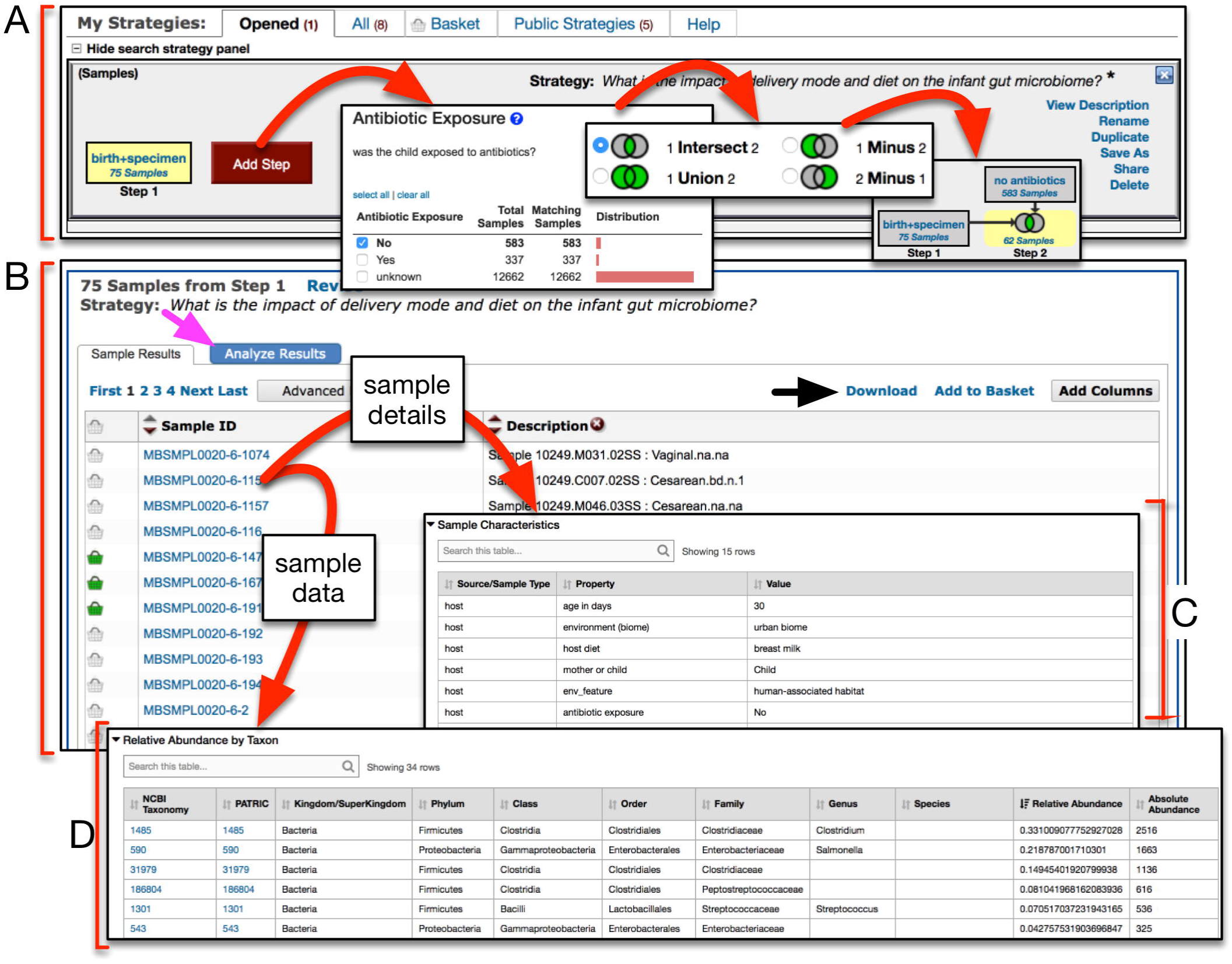
Screenshot of the query results page. (A) The strategy panel provides users with an interface to name, share and expand on their initial query, thereby constructing *in silico* experiments. Query results are shown as ‘Step 1’, and additional queries can be added as additional steps. (B) The sample results panel shows all samples matching the search strategy, which can be downloaded (black arrow). Users can visualize and statistically analyze their query results by accessing a suite of interactive web apps (magenta arrow). (C-D) Details and data for individual samples can be viewed by clicking on the sample identifier.

## DATA VISUALIZATION AND ANALYSIS

Clicking on the ‘Analyze Results’ tab of the results page (**Figure 4B, magenta arrow**) reveals a suite of interactive web apps which allow users to visualize and statistically analyze the samples returned from any query, directly in the browser window **(Figure 5A)**. All apps were built using the Shiny framework (28, 29) which allows the development of web-based applications using the R programming environment (26). Our Shiny Apps follow four common guiding principles: first, all data and sample details returned by a query strategy are passed to the app so that users can explore their data in the context of the experimental covariates. For example, graphs can be faceted and colored to reveal how factors such as diet, specimen type, or disease status are associated with shifts in microbial community diversity or composition. Second, all apps adhere to the ‘grammar of graphics’ and were generated using ggplot2 (25). Third, when appropriate, non-parametric statistical analyses (Wilcoxon or Kruskal-Wallis rank sum test) are automatically computed after faceting and test results are displayed on the graphic. Fourth, any graphic produced and the underlying data can be downloaded directly from within the app. Currently, five apps are available on MicrobiomeDB, and these are described in more detail below.

**Figure 5:**
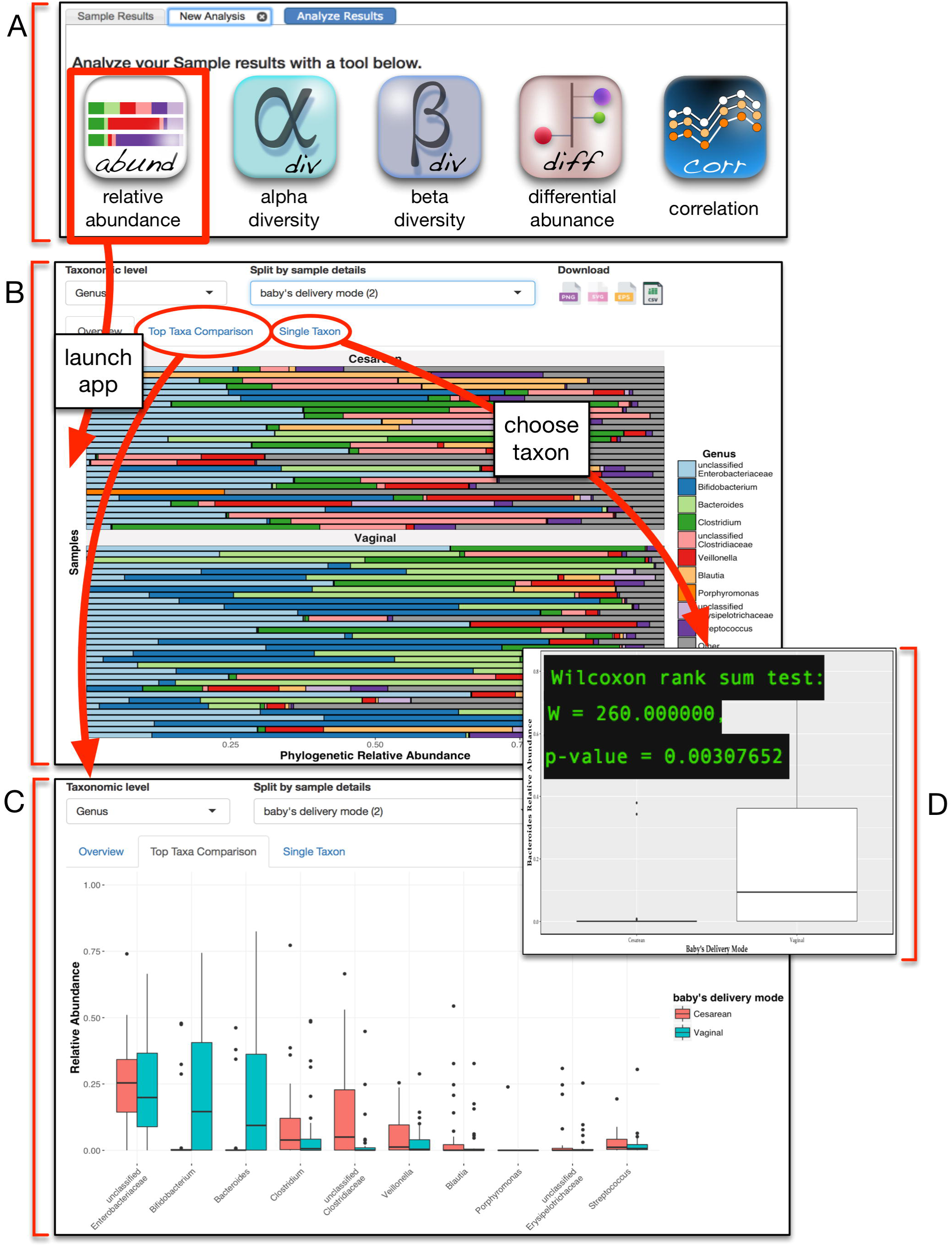
Screenshot of the relative abundance app. (A) The analysis tab of the results page provides access to a suite of interactive web apps for visualization and analysis of microbial community diversity and composition. (B) Selecting the relative abundance app displays a horizontal stacked bar chart of the top ten most abundant taxa. Users can customize this graphic by selecting taxonomic level and sample details to partition the samples into groups. (C) Navigating to the ‘top taxa comparison’ tab of the app displays this data as a box-and-whisker plot that allows the same customization as the stacked bar chart. (D) Double clicking on any single taxon in panel B, or navigating to the ‘single taxon’ tab of the app and entering a taxon of interest, displays a graph of that taxon with statistical analysis comparing the relative abundance between the user-defined group(s).

### Relative abundance app

Relative abundance of taxa is pre-calculated for each sample during the data loading workflow (**Figure 1)**. When the relative abundance app is launched, a horizontal stacked bar chart is created from the top 10 most abundant taxa present (by median relative abundance) across all the samples returned by the query, and the relative abundance for all remaining taxa is binned together and displayed as an 11^th^ group termed ‘other’ (**Figure 5B**). A drop down menu is available to change the taxonomic rank, or to partition the graph based on any available covariates for the displayed samples, producing an updated graphic with each new user input. Selecting the ‘Top Taxa Comparison’ tab of this app opens a new graphic that displays the top 10 taxa as box-and-whisker plots, with one box per covariate (**Figure 5C**). Finally, a third tab of this app, ‘Single Taxon’, provides users with a searchable list of all taxa present in the samples. Selecting a taxon from this list produces a box-and-whisker plot for only that taxon, and calculates significance **(Figure 5D)**.

### Diversity apps

Unlike relative abundance data, diversity metrics are not pre-calculated at the time of data loading. Instead, when users launch the alpha diversity app, the PhyloSeq package (26) is used to *de novo* calculate Shannon, Simpson, Chao1, ACE and Fisher diversity metrics, which are then displayed as either a dot or a box plot **(Figure 6B)**. Clicking on the ‘Explore Sample Details’ tab of the app **(Figure 6B, arrow)** allows users to facet the plot based on one or more experimental variables. Similarly, when users launch the beta diversity app, Bray-Curtis, Jansen-Shannon divergence, Jaccard, Kulczynski, Canberra, Horn, and Mountford metrics are used to calculate dissimilarity between samples, which is then used to ordinate the samples as points on a two-dimensional plot, where point color and shape can be mapped to sample details **(Figure 6C)**.

**Figure 6:**
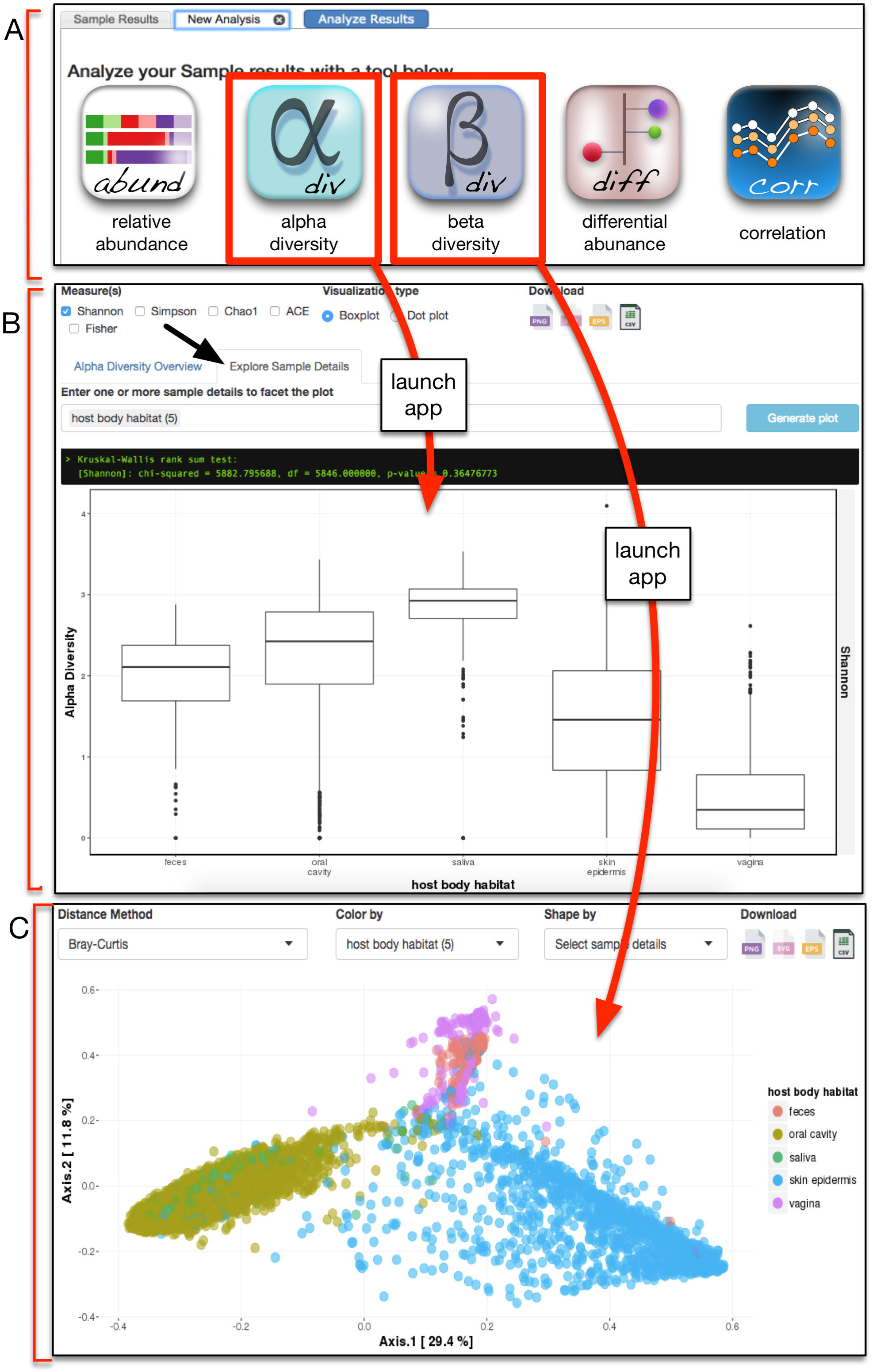
Screenshot of the alpha and beta diversity apps. (A) The analysis tab of the results page provides access to a suite of interactive web apps for visualization and analysis of microbial community diversity and composition. (B) Launching the alpha diversity app allows users to choose between Shannon, Simpson, Chao1, ACE and Fisher diversity metrics, and to display the data as either a box plot or a dot plot. Selecting the ‘Explore Sample Details’ tab of the alpha diversity app (black arrow) lets users facet the plot using any available sample details. (C) Similarly, launching the beta diversity app presents users with drop down menus to choose distance method, and to color and shape points based on sample details. In the example shown in panels B and C, alpha and beta diversity were analyzed by host body habitat for over 6000 samples from the Human Microbiome Project.

### Differential abundance and correlation apps

Launching the differential abundance app **(Figure 7)** allows users to formally test for differentially abundant taxa between any pairwise comparison of sample details using DESeq2 (27). The user selects the experimental variable of interest from a drop down menu (e.g. baby’s delivery mode) and is presented with two additional drop down menus that show all values associated with the selected term. Choosing the pairwise comparison of values (e.g. cesarean versus vaginal) initiates the differential abundance analysis. Results are displayed as a ‘lollipop’ chart where each lollipop represents a differentially abundant taxon, its direction indicates the phenotype association, length indicates fold change, the color indicates phylum, and the size signifies statistical significance **(Figure 7B).** Moving the cursor over any taxon displays a box-and-whisker graph and statistics for that taxon **(Figure 7C)**. Rather than specifically testing for differentially abundant taxa, users may want to explore associations between taxa and sample details more broadly. To address this need, the correlation app **(Figure 7A)** displays the Spearman’s rank correlation between sample details and taxa, and also allows users to also view correlations between different sample details. The result is shown as a dot plot where the colors indicate the Spearman’s rho (blue for positive correlation and red for negative correlation) and the size signifies statistical significance. A searchable table is also displayed under the chart where the user can see all the correlations in a table structured way.

**Figure 7:**
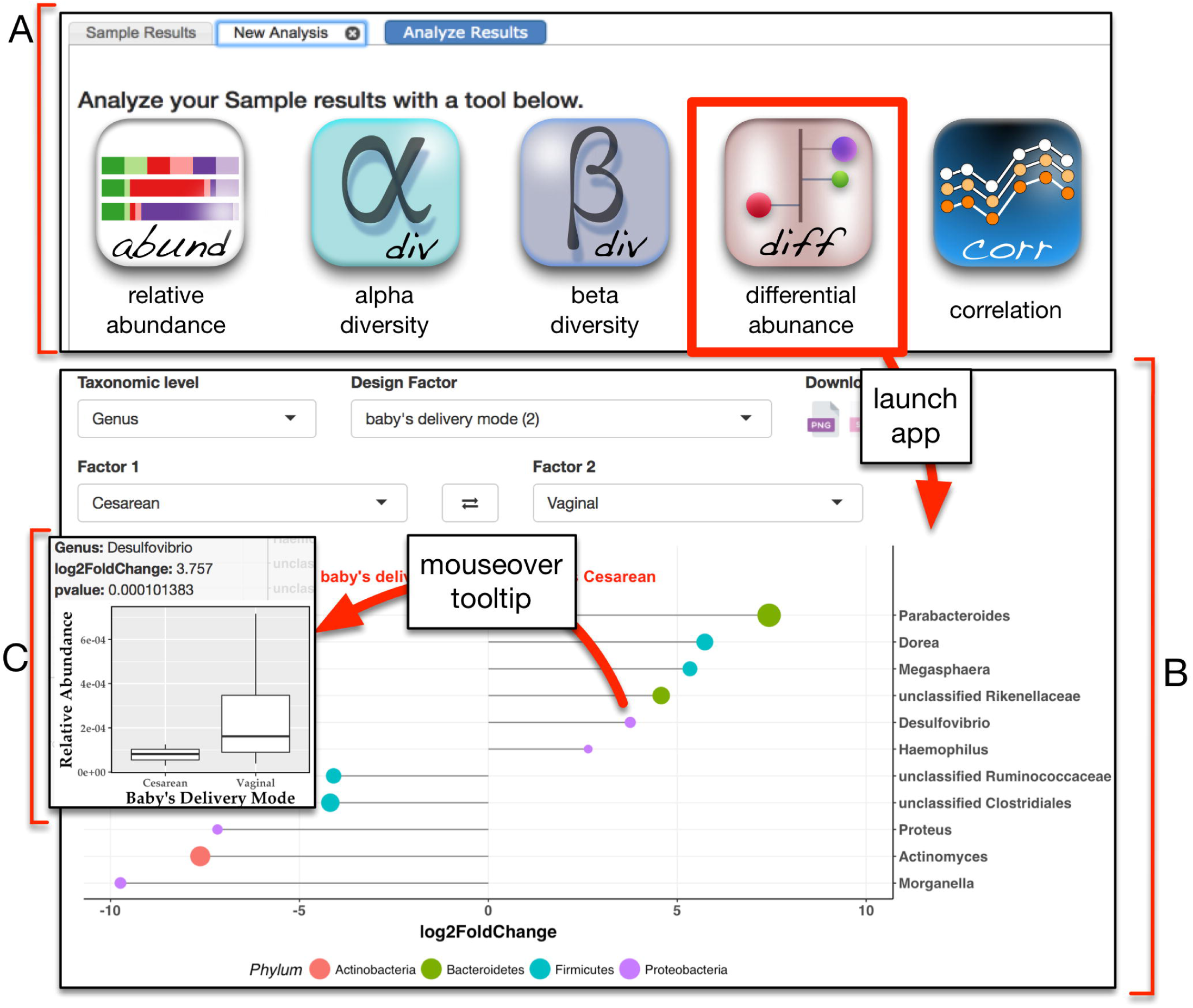
Screenshot of the differential abundance app. (A) The analysis tab of the results page provides access to a suite of interactive web apps for visualization and analysis of microbial community diversity and composition. (B) Selecting the differential abundance app presents users with several drop down menus to customize their analysis. After choosing the taxonomic level, the design factor, and the pairwise comparison of interest, DESeq2 is run to identify differentially abundant taxa. Results are displayed as a ‘lollipop’ chart where color indicates phylum, length of the lollipop indicates log2 fold change (X-axis), and size of the lollipop reflects statistical significance. (C) Moving the cursor over any lollipop displays a plot of relative abundance with statistics for that taxon.

## ADDITIONAL FEATURES

### Favorites and Basket

Through the favorites tool, users can bookmark samples of interest for convenient access later. Adding or removing a sample to the favorites can be done by clicking on the favorites icon (star) present at the top of each sample record page (**Figure 4C and 2D**). Samples in the favorites page can be assigned to user-defined projects and free text can be added to describe each sample. Additional flexibility for dealing with custom sample lists is provided by the basket tool, which allows a user to compile and save a set of samples that can be added to a search strategy and thereby incorporated with any other samples from MicrobiomeDB. Samples are added to the basket by clicking on the basket icon next to the sample ID on the search results page **(Figure 4B)** or at the top of a sample record page (**Figure 2C and 2D**). Access to favorites or the basket page is achieved via links in the menu bar at the top of the homepage.

## FUTURE DIRECTIONS

Future development efforts for MicrobiomeDB will focus on three objectives: First, the number of available datasets will be dramatically expanded. Although we eventually expect to load all datasets available through the QIITA portal (https://qiita.ucsd.edu/) (7, 8), our initial focus will be on publically available microbiome datasets generated from gastrointestinal diseases in humans and animals, including inflammatory bowel disease, infection and malnutrition. As the number of studies loaded into MicrobiomeDB grows, we anticipate that the custom application ontology used on the site to describe samples (available for download through our datasets page) will become a valuable resource for microbiome researchers. Second, the data loading workflow will be designed to handle not only.biom files, but also raw 16S rRNA gene sequence data, ensuring standardized data processing as part of the workflow. Finally, data visualization and analysis apps will continue to be refined and new apps added to the analysis suite, including new methods to find taxa associated with continuous experimental variables, such as age and weight (https://doi.org/10.1101/099457). It is common for microbiome experiments to include other assays, such as metabolomics or transcriptional profiling, and the extensibility of our data loading workflow and the EuPathDB toolkit provide an opportunity to incorporate diverse data types. Taken together, these results constitute a first step toward a full-featured, open-source web platform for a systems biology view of microbial communities.

---

**Table S1:** Datasets currently available in MicrobiomeDB. Microbial census data (.biom) and sample details for 14179 samples from five different studies were downloaded from QIITA (https://qiita.ucsd.edu/) and loaded into MicrobiomeDB. The study title and ID from QIITA are shown.

